# SARS-CoV-2 Intra-host Variation Shows Evidence of Transmission and Convergent Evolution in a University Surveillance Cohort

**DOI:** 10.64898/2025.12.12.694004

**Authors:** Léa Cavalli, Bradford P. Taylor, Beau Schaeffer, Jacquelyn Turcinovic, John H. Connor, William P. Hanage

## Abstract

Monitoring and understanding the transmission and evolution of SARS-CoV-2 remains a significant pub-lic health priority. Within-host genetic variation provides insight into viral evolution during infection and may help infer transmission events. In this study, we analyzed intrahost variation in SARS-CoV-2 genome sequences from Boston University’s testing mandate. Focusing on intrahost single nucleotide variants (iSNVs), we inferred transmission events and assessed the selective forces shaping within-host viral evolution. To minimize false-positive iSNVs resulting from systematic biases, we implemented stringent data filtering and developed a heuristic to exclude contamination-derived artifacts arising from batched sequencing. We find that intrahost variation is limited and infrequently transmitted during acute infections, suggesting that shared iSNVs serve as highly specific but insensitive markers of transmission. We also observed incomplete purifying selection shaping within-host diversity, with the loci most affected changing among variants of concern. Finally, we identified a highly recurrent iSNV (G11083T) which may represent a site of positive selection. Our results highlight that within host variation provides insight towards within host pathogen evolution, in spite of a limited use towards genomic epidemiology.

**IMPACT STATEMENT:** SARS-CoV-2 is the most extensively sequenced pathogen to date, yet much of its genomic data remains underutilized. Intrahost variation, in particular, is less studied than consensus-level variation, partly because most datasets lack technical sequencing replicates to control false-positive signals. Using genomic data from a university testing mandate and applying rigorous filtering to systematically minimize false-positive iSNVs in a data-driven manner, we obtain insights into SARS-CoV-2 evolution and transmission from intrahost variation. Our work underscores the potential to use existing, large-scale datasets to better understand pathogen evolution in situ.

**DATA SUMMARY:** All sequence data have been deposited in the Sequence Read Archive (SRA) of the national center for biotechnology information (NCBI), under the project accession number PRJNA892225. All code is open-access and available in GitHub at: https://github.com/Leacavalli/Sars-cov-2-Intrahost-Variation. Any additional supporting data has been provided within the article.

## INTRODUCTION

Over the course of the COVID-19 pandemic, genomic surveillance has documented the evolution of SARS-CoV-2 in unprecedented fashion. The great majority of genomic data generated consist of consensus sequences, which have been instrumental for identifying single nucleotide polymorphisms (SNPs) associated with specific phenotypic traits such as replication rate, transmissibility, virulence, and immune evasion, and that define Variants of Concern (VoCs e.g. Alpha, Delta, Omicron) [1, 2, 3, 4] These variants have exhibited unusual amounts of divergence from currently circulating strains and are widely considered to have emerged from persistent infections in immunocompromised hosts. [5] Multiple studies of such infections have examined subconsensus alleles, which reflect intra-host variation arising over the course of infection, and shown how mutations arise and sweep to fixation. In contrast, fewer studies have examined intra-host variation during acute infections, even though it may contribute to the diversity observed within circulating variants.

Intra-host single nucleotide variant (iSNVs) can, in theory, be used to infer both the selection pressures acting during infection and the transmission patterns of intra-host variation. [6, 7, 8, 9] In practice, however, iSNVs are particularly susceptible to sequencing artifacts and well-to-well contamination because they represent only a minority of reads at any given site. Technical sequencing replicates, used to identify and remove false-positive signals, have shown that many putative iSNVs fall into this category.[7, 8, 10] However, most sequence data are generated through routine genomic surveillance, which does not include such resource-intensive experimental controls. While filtering sequencing reads and variants based on quality control thresholds can help minimize false positive iSNVs arising from sequencing artifacts, there have been limited efforts to develop a standardized data-driven method to address false positives arising from contamination. [11] Instead, potential contamination is often assessed on a case-by-case basis for iSNVs of interest, for example by checking for batch effects or flagging iSNVs that match lineage-defining mutations. [12] The development of a standardized approach to systematically detect and mitigate contamination-related false positives would allow analysis of subconsensus variation in large datasets without technical sequencing replicates.

In this study, we analyzed SARS-CoV-2 genome sequence data from a sample collected through Boston University’s testing mandate from January 2021 to April 2022, and investigated SARS-CoV-2 intra-host variation in viral genomes that were classified into the Delta, BA.1 and BA.2 VoCs. To overcome the absence of direct experimental controls, we applied stringent filtering to minimize sequencing bias, using batch information to systematically remove false-positive iSNVs resulting from contamination. This enabled us to identify potential transmission pairs and investigate the evolutionary forces acting on SARS-CoV-2 within individual hosts, yielding observations consistent with those reported in carefully controlled studies.

## METHODS

### Data Collection and Sequencing

BU collected anterior nares swabs from students, staff, and faculty on a weekly basis as part of its university-wide SARS-CoV-2 surveillance testing program in response to the pandemic. [13] After diagnostic testing, positive samples were used for whole viral genome sequencing. Total RNA was prepared from 300ul of starting sample using the Zymo Quick-RNA Viral 96 Kit according to the manufacturer’s instructions. SARS-CoV-2 genomes were amplified from the total RNA preparation using the Artic protocol and recommended primer sets (versions 3-4). Following rtPCR, amplicon pools were combined, barcoded and sequenced using Illumina SBS approaches. Sequencing was performed by BU’s Microarray and Sequencing Resource.

### Bioinformatic processing

Reads were trimmed with Trimmomatic (v.0.39), and successful trimming was verified with FastQC. [14, 15] Trimmed reads were aligned to the Wuhan-Hu-1 reference genome (MN908947.3) using BWA (v0.7.17-r1188). [16, 17] Primer sequences were soft-clipped with iVAR (v.1.4.4) before variant calling with Lofreq (v. 2.1.5). [18, 19] In Lofreq, we excluded variants with alternative allele frequency (AAF) lower than 0.01, coverage lower than 100, quality lower than 30 and significant strand bias (SB p-value <0.01 with fdr multiple testing correction). [19] Functional annotation was performed with SnpEff (v 5.2). [20] We used bcftools consensus (v.1.22) to generate consensus sequences from major variants (AAF ≥ 0.5), setting low coverage positions (<100) set to ‘N’. [21] Pangolin was run in accurate (UShER) mode (v 4.3) for lineage assignment. [22] These processing steps were implemented in a custom Nextflow (v 23.10.0.5889) pipeline (Figure S1). [23] After running the pipeline on individual samples, Mafft (v7.520) was run to create a multiple sequence alignment (MSA), used to infer a maximum likelihood (ML) phylogeny with RAxML (v8.2.12). [24, 25] Inference was performed under the GTR + Γ model, and the resulting tree was rooted with the Wuhan-Hu-1 reference. Upon confirming temporal structure with a root-to-tip regression generated with Treetime (v0.11.4), we used IQtree to construct a time-calibrated tree (v 2.2.5). [26, 27] All code, along with a flowchart detailing the bioinformatic methods and a table of parameter values, is available on GitHub at: https://github.com/Leacavalli/Sars-cov-2-Intrahost-Variation.

### Minimizing False Positive iSNVs from Systematic Bias

The data filtering steps described below are illustrated in a flowchart in fig. S2. They were implemented in R (v.4.3.1) [28], and the code is available in GitHub.

#### Exclusion of low quality isolates

The initial dataset consisted of 4,295 SARS-CoV-2 isolates. We excluded 390 isolates with missing sequencing batch information, as contamination could not have been confidently identified in these cases. In addition, we removed 619 isolates with low viral loads (Ct values ≥ 32), and 348 isolates with *>* 10% missing bases (`N` or gaps) in their consensus sequences, 103 isolates from pangolin lineages corresponding to VoCs other than Delta, BA.1, or BA.2 (*n* = 100 *Alpha, n* = 1 *Iota, n* = 39 *B.1.1.529, n* = 3 *B.1, n* = 4 *B.1.1*), and 68 phylogenetic outliers, defined as isolates that had average pairwise genetic distances to other isolates within their VoC more than three standard deviations from the mean, as well as outliers identified by TreeTime. Our final dataset consisted of 2,770 SARS-CoV-2 genomes from *Delta* (*n* = 777), *BA.1* (*n* = 1,199), and *BA.2* (*n* = 794) isolates. The quality characteristics of the final samples included in our study are summarized in Figure S3. These yielded 138,259 fixed SNPs with an alternative allele frequency (AAF) of 0.95 or higher, and 212,345 potential iSNVs, with AAFs between 0.01 and 0.95.

#### Exclusion of false positive iSNVs from cross-contamination

The most recurring iSNVs commonly exhibited patterns consistent with well-to-well contamination across sequencing batches (See Supplementary text). In response, we developed and applied a heuristic to systematically identify putative false-positive iSNVs on the basis of sequencing batches, AAFs, and lineage-defining mutations. Following the identification of putative contamination (9,497 iSNVs), we retained either the majority allele as a fixed SNP (n=2,815) or the reference allele (n=6,682). We detail our heuristic and its limitations in the supplementary text.

#### Exclusion of Residual Sequencing or Contamination Artifacts

To remove iSNVs arising from residual sequencing artifacts, we excluded all iSNVs with AAF < 0.03, a threshold shown in replicate-based studies to provide optimal sensitivity and specificity. [6, 29] Before lowering the AAF threshold, however, we used low-frequency iSNVs to identify additional false positives not removed by AAF-based filtering alone (described in the supplementary text). We removed any iSNVs recurring at minor allele frequency (0.01 ≤ AF *<* 0.05) in isolates across five or more sequencing batches (n = 35,708), or across two or more isolates within the same batch (n = 122,640). In addition, iSNVs that recurred within a batch more than twice at major allele frequency (AF ≥ 0.5), including at least one instance at fixed frequency (AF ≥ 0.95) (n = 186), were reclassified as fixed SNPs. All such patterns are more likely to be artifacts rather than convergent evolution or transmission clusters, though this filtering is conservative. Finally, we restored any C22792T iSNV to a fixed AAF, since it is a known artifact associated with primer binding issues (Figure S6). [30]

### Transmission Analyses

The consensus SNP differences used to infer potential transmission pairs were estimated by multiplying the reference genome length (29,903 bp) by patristic distances, calculated from the phylogenetic tree using the `cophenetic.phylo` function from the `ape` R package. [31] Although correlated with SNP counts, this measure provides a more accurate estimate of genetic differences between samples (Figure S7).[32, 33] Links between likely transmission pairs were mapped onto the time-calibrated phylogeny using the R packages `ggraph` and `ggtree`.[34, 35]

### Selection Analyses

To identify evidence of selection dN/dS ratios were computed with the `dNdScv` R package (v. 0.0.1.0) [36] using the default substitution model and the SARS-CoV-2 reference file developed by Tonkin-Hill, Martincorena, Amato, et al., (2021). To estimate the dN/dS at the within host level, all iSNVs (0.03 ≤ AF *<* 0.95) were used. In contrast, at the population level, using all fixed SNPs (AF ≥ 0.95) across isolates can lead to inflated dN/dS values with this package, as lineage-defining mutations carried by many samples (i.e., homoplasies) are counted multiple times. (Figure S8) To control for this, we counted each fixed SNP only as many times as it independently evolved, using a mutation-annotated phylogeny constructed with Usher and MatUtils.[37, 38]

We note that although the dN/dS ratio is increasingly frequently applied to within-host variation, it was originally designed to assess fixed mutations in populations that have diverged over extended evolutionary timescales. This metric is thus inherently time-dependent and its interpretation over the much shorter timescales of infections requires caution.[39] Specifically, dN/dS can be artificially inflated for recent changes, as purifying selection may not have had sufficient time to eliminate new deleterious nonsynonymous mutations arising through the random mutation process. In addition, dN/dS may not be sufficiently sensitive to detect recent positive selection, particularly in contexts where purifying selection predominates, as is the case for SARS-CoV-2 within hosts.

## RESULTS AND DISCUSSION

### Characteristics of true intrahost variation

6,128 iSNVs were found in the complete dataset of 2,770 samples following removal of putative false-positive iSNVs from sequencing artifacts and potential contamination as detailed in the Methods. The characteristics of these iSNVs are reported in Table 1. Briefly, around half of all samples contained iSNVs, a proportion that did not vary greatly among the different VoCs. While the median number of iSNVs was 0 for all variants, the mean and 3rd quantile were both 2 (ranging from 0-54 in Delta, 0-44 in BA.1 and 0-25 in BA.2, See Figure S9a), reflecting a similar distribution to those previously reported. [6, 8, 12] This is consistent with limited diversity during the acute phase of infection, and contrasts with the accumulation of mutations in persistent infections. [5] The small number of samples with large numbers of iSNVs underline that such infections are rare. Most of these samples appeared on different sequencing plates and the iSNVs they carried did not match lineage-defining SNPs, ruling out residual contamination or mixed infection. However, the iSNVs were largely present at very low frequencies, a pattern more consistent with sequencing artifacts than with persistent infection or hypermutation, though this cannot be determined with certainty without technical replicates.

**Table 1.** Characteristics of iSNVs and Fixed SNPs across VoCs.

|  | Delta<br>(n=777) | BA.1<br>(n=1,199) | BA.2<br>(n=794) | Overall<br>(n=2,770) |
| --- | --- | --- | --- | --- |
| **iSNVs (Real)** |  |  |  |  |
| Number of Samples with iSNVs (%) | 388 (49.9%) | 649 (54.1%) | 425 (53.5%) | 1,462 (15.3%) |
| Total iSNVs (median; mean) | 1,758 | 2,636 | 1,734 | 6,128 |
| (median; mean) | (0; 2.3) | (0; 2.2) | (0; 2.2) | (0; 2.2) |
| Non-coding | 83 | 74 | 35 | 192 |
| Synonymous (S) | 468 | 851 | 560 | 1,879 |
| Non-synonymous (N) |  |  |  |  |
| Missense ($N_M$) | 1,149 | 1,656 | 1,082 | 3,887 |
| Nonsense ($N_N$) | 58 | 55 | 57 | 170 |
| N/S | 2.46 | 1.95 | 1.93 | 2.07 |
| $d_N/d_S$ (95% CI) | 0.81 (0.72–0.92) | 0.83 (0.76–0.91) | 0.80 (0.72–0.90) | 0.82 (0.77–0.87) |
| **Fixed SNPs (fSNPs)** |  |  |  |  |
| Total | 28,416 | 64,309 | 48,601 | 141,326 |
| Total homoplastic sites* | 1,033 | 345 | 318 | 1,753** |
| Non-coding | 36 | 13 | 3 | 54** |
| Synonymous (S) | 396 | 144 | 143 | 683** |
| Non-synonymous (N) |  |  |  |  |
| Missense ($N_M$) | 596 | 187 | 172 | 955** |
| Nonsense ($N_N$) | 5 | 1 | 0 | 6** |
| N/S | 1.52 | 1.31 | 1.20 | 1.45 |
| $d_N/d_S$ (95% CI) | 0.49 (0.43–0.57) | 0.48 (0.38–0.60) | 0.50 (0.40–0.64) | 0.51 (0.45–0.56) |

\* Each fixed SNP is counted only as many times as it independently arose, such that convergent (homoplastic) events are counted once per independent origin (see Methods).
\*\* These numbers may be greater than the sum of corresponding values across VoCs due to the fixed mutations present on nodes leading to each VoC.

### Most Shared iSNVs Are Attributable Recurrent Mutation

The great majority of iSNVs occurred in a single sample, but about 5% were found in two or more, potentially reflecting transmission or convergent evolution (Figure S9b). One particularly recurring iSNV (G11083T) was found in 20 separate samples (17 of them Delta, 3 BA.1) and is discussed in detail below. Of the 1,056 pairs that shared iSNVs, most shared a single iSNV (n= 1,039), some shared two iSNVs (n=14), and 3 pairs shared three iSNVs. Among pairs that shared multiple iSNVs, none corresponded to mutations that were fixed at consensus in any lineage, indicating that these were not mixed infections or residual contamination. In addition, many pairs that shared iSNVs were distantly related on the phylogeny, even being in different VoCs (Figure 1). These findings are consistent with shared iSNVs arising from recurrent mutation, rather than co-infections or the transmission of intra-host variation. [8] Recurrent mutations may be the result of biases in the mutation spectra, or convergent evolution, as further discussed below.

**Figure 1.**
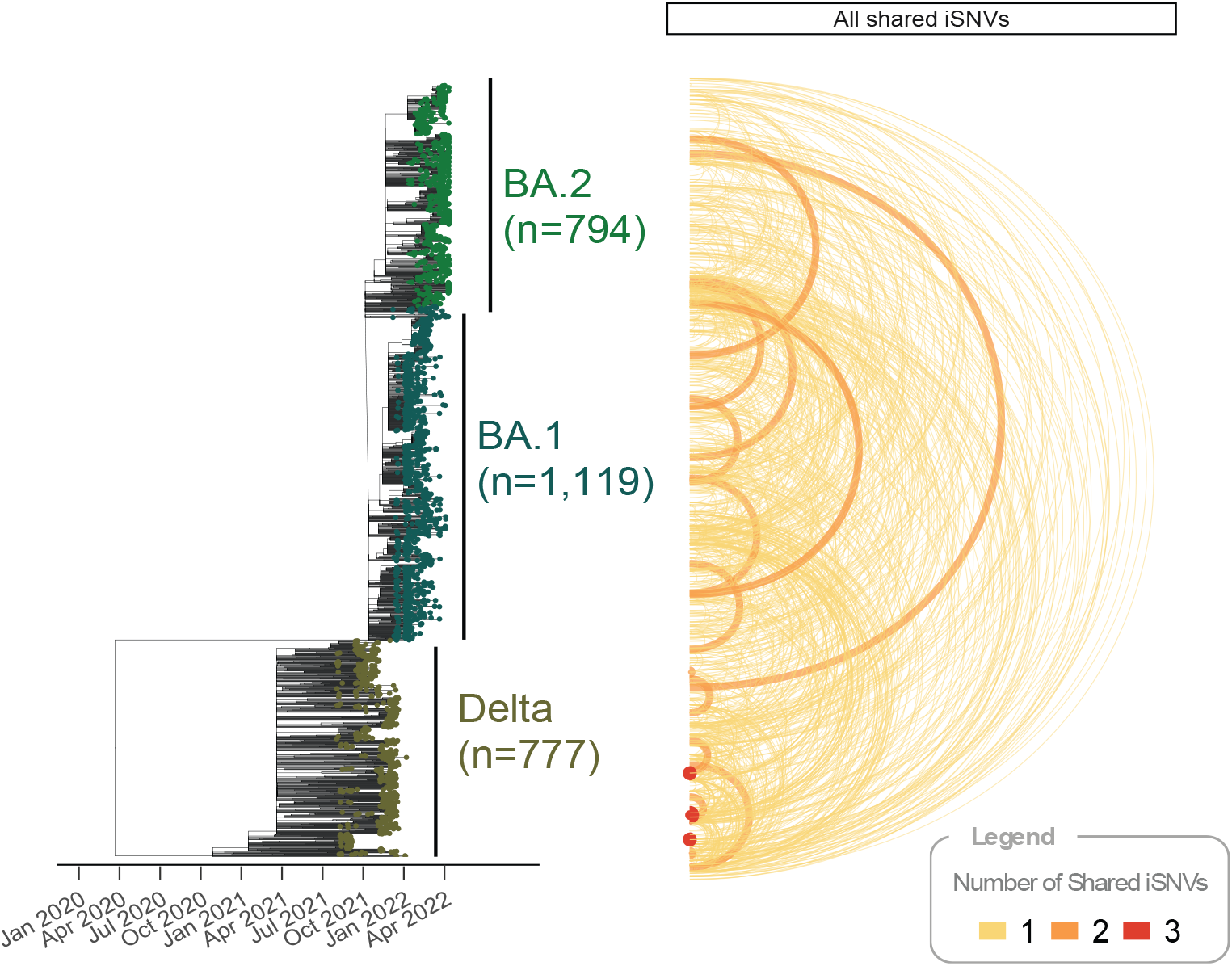
Time-calibrated phylogeny with links indicating shared variation. Link width and color indicate the number of shared iSNVs; link height reflects phylogenetic distance between tips.

### Inferring Transmission from Intrahost Variation

#### Identification of Potential Transmission Pairs

Shared iSNVs have been suggested as a marker of transmission, the rationale being that it is extremely unlikely two closely related infections would have mutations at the same sites by chance. In the case of SARS-CoV-2, limited genetic diversity is a challenge when resolving transmission, and shared iSNVs could supply an additional level of resolution. [33] Therefore, we first identified potential transmission events as sample pairs collected within 10 days of each other and differing by two or fewer consensus SNPs. The characteristics of potential transmission events are shown in Table 2. We identified a total of 3,919 potential transmission events, often grouped together into clusters containing multiple potential sources of infection (Figure 2). Indeed, 136 potential transmission events involving 122 samples were identified among Delta isolates. Because a maximum of only 121 transmission events is possible for 122 samples (assuming no superinfections), this indicates that 15 of the inferred events are redundant, arising from the inability to identify the true transmission source among multiple pairs of closely related samples. In BA.1, 3,358 potential transmission events were identified from 652 samples, yielding 2,707 redundant links based on consensus variation. In BA.2, 1,304 potential transmission events were identified from 338 samples, resulting in 966 redundant links. Therefore, we interpreted the size of these clusters as the maximum number of potential transmission events captured in our dataset, which allowed us to estimate a maximum number of locally acquired infections (i.e., within the BU community) and a minimum number of importations from outside of the university community (see Table 2). Clusters were larger for BA.1 and BA.2, consistent with the high transmissibility of these variants. [40, 41] Shifts in network structure and social connectivity over time also will have altered opportunities for introduction and transmission. For example, the Delta variant exhibited the highest proportion of importations, and its peak prevalence in our dataset occurred in late November–early December 2021 (Figure S3)), a period overlapping with major holidays when many students were likely traveling to and from campus.

**Table 2.** Summary of Potential Transmission Events across VoCs.

|  | Delta | BA.1 | BA.2 | Overall |
| --- | --- | --- | --- | --- |
| Total isolates ($N$) | 777 | 1,199 | 794 | 2,770 |
| **Potential transmission events** |  |  |  |  |
| Overall | 136 | 3,358 | 425 | 3,919 |
| With shared iSNVs | 3 (2.2%) | 4 (0.12%) | 2 (0.15%) | 9 (0.23%) |
| With fixation events | 1 (2.6%) | 3 (0.88%) | 0 (0%) | 4 (0.10%) |
| Number of transmission clusters($N_c$) | 39 | 62 | 48 | 149 |
| Total isolates in transmission clusters($N_t$) | 122 | 652 | 339 | 1,113 |
| Estimated locally-acquired infections ($N_l = N_t - N_c$) | 83 (11%) | 590 (49%) | 291 (37%) | 964 (35%) |
| Estimated imported infections ($N - N_l$) | 694 (89%) | 609 (51%) | 503 (63%) | 1,746 (63%) |

**Figure 2.**
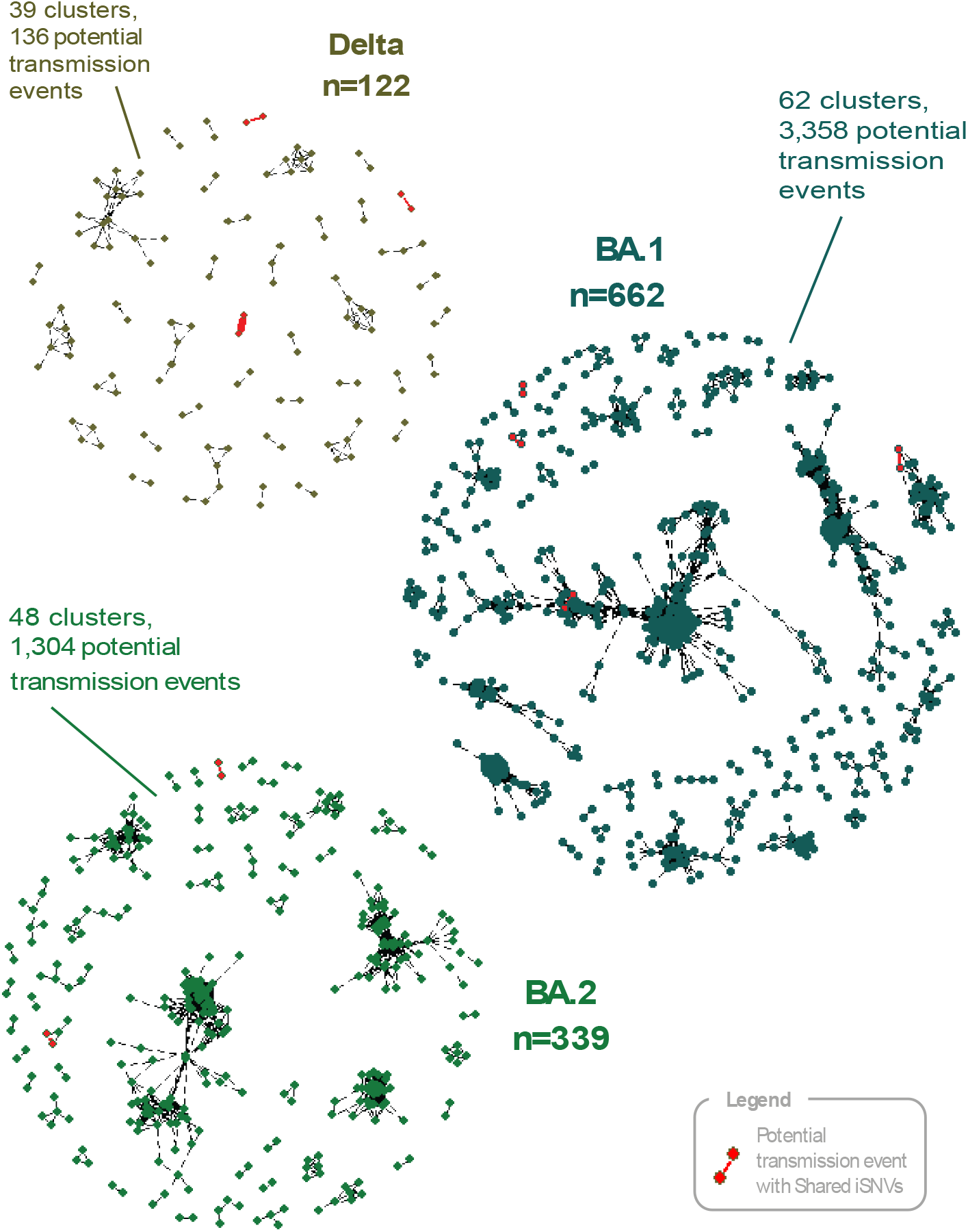
Network of Potential Transmission Events Among Delta, BA.1 and BA.2 Isolates. Each link represents a potential transmission pair, defined as a pair of isolates collected within 10 days of each other and with an estimated consensus SNP distance of two or fewer. Black links indicate sample pairs that do not share iSNVs, while red links indicate pairs that share iSNVs; the width of each link corresponds to the number of shared iSNVs. Among Delta samples, 122 were involved in 136 potential transmission events forming 39 clusters. Among BA.1 samples, 662 were involved in 3,358 potential transmission events forming 62 clusters. Among BA.2 samples, 339 were involved in 1,304 potential transmission events forming 48 clusters.

#### Shared iSNVs may be a highly specific but insensitive signature of transmission

Of the 3,919 potential transmission events identified as above, only 9 were found to share iSNVs (see Table 2). With one exception in all cases the samples shared a single iSNV. The exception of one pair of Delta isolates that shared three iSNVs (C25160T, G27390T, and G29737T). We note that, among the other two isolate pairs sharing three iSNVs, one pair - collected 6 days apart, differing by three consensus SNPs and not sequenced on the same plate - may represent a transmission event missed by our stringent criteria, while the other - collected one day apart, differing by nine consensus SNPs, and sequenced on the same plate – may be a case of mixed infection or residual contamination. These findings are consistent with others that imply an extremely tight bottleneck in the majority of transmission events such that the co-transmission of subconsensus variants is rare. [6, 42] Although uncommon, the detection of shared iSNVs in potential transmission pairs indicates that multiple subconsensus alleles can rarely be transmitted together from one individual to another. [6] Therefore, shared variation likely lacks sensitivity for broadly inferring transmission events, since it does not manifest in many closely related pairs. However, when present, it may serve as a highly specific indicator of transmission, useful for disentangling multiple potential transmission sources. For instance, of the nine potential transmission events with shared iSNVs, four were involved in clusters with several possible sources, such that the shared variation could identify the most plausible transmission link.

#### Fixation of iSNVs Through Transmission is Rare

Shared iSNVs are not only a signature of transmission; all the extant genetic diversity in the SARS-CoV-2 population started as an iSNV that fixed in the next infection, either by selection or drift. Therefore, we also investigated whether we could identify iSNVs that appeared to move from subconsensus to fixation upon transmission. We found 4 potential transmission pairs where an iSNV in one isolate was a fixed SNP (AF ≥ 0.95) in the other: 1 of Delta isolates and 3 BA.1, and no BA.2 (Figure 1). All such sample pairs shared a single mutation, except one pair of BA.1 isolates that shared mutations at two positions. Overall, the iSNVs linked to possible fixation events described here and the shared iSNVs among transmission pairs described above both exhibited relatively high AAFs (Figure S10), suggesting that iSNVs must reach sufficiently high frequencies in the donor in order to be transmitted.

The limited number of potential co-transmission and de novo iSNV fixation events observed in our study could be a consequence of our conservative filtering strategy, which prioritizes sensitivity over specificity, and consequently removes some true positive iSNVs. Indeed, some true iSNVs involved in co-transmission or fixation events may be classified as contamination according to our heuristic if both involved samples were present on the same sequencing plate, which is likely since transmission pairs are typically sampled around the same time. This methodological limitation may also contribute to the lack of co-infections observed in our study compared to previously reported rates (0.2–0.6%), although these estimates may have themselves been biased by contamination. [43]

Overall, the limited intra-host variation observed in our study, together with its very rare transmission, illustrates the impact of the transmission bottleneck on the virus’ effective population size and its mutation supply. This limits SARS-CoV-2’s capacity to explore evolutionary space over acute infections, which is consistent with prolonged chronic infections being the source of new VoCs with substantial immune escape, since they provide both a larger effective population size and more time to evolve antigenic shift. [44] Acute infections may still, however, remain a source of the antigenic drift that drives the more subtle variation observed among sublineages of circulating VoCs.

### Investigation of Within-Host Selection Pressures

#### Incomplete Purifying Selection Within Hosts

Overall, we identified 5,936 iSNVs in coding regions across all samples, including 4,057 non-synonymous and 1,879 synonymous mutations (Table 1). The ratio of nonsynonymous to synonymous substitution rates (dN/dS) among iSNVs was 0.81 (95% CI: 0.76-0.86), indicative of mild purifying selection, as previously reported. [7, 8, 12, 45, 29] In contrast, the dN/dS of fixed SNPs was 0.51 (0.45-0.56), indicating stronger purifying selection among fixed SNPs compared to iSNVs (Figure S8). These observations are aligned with the two-step fitness selection process proposed by Li et al. (2022), whereby non-synonymous iSNVs occur at a higher frequency within hosts compared to the population level because purifying selection over the course of infection is unable to fully eliminate new deleterious mutations (i.e., incomplete purifying selection). [7]

#### Intrahost Selection Processes across VoCs

Although we found no evidence for overall differences in the dN/dS rates of iSNVs across VoCs (Table 1), the relative frequencies of missense, nonsense, and synonymous mutations among iSNVs and the corresponding dN/dS rates suggested cross-VoC variation in selection on specific genomic regions, while others were in common between Delta and BA.2 (Figure 3A).

**Figure 3.**
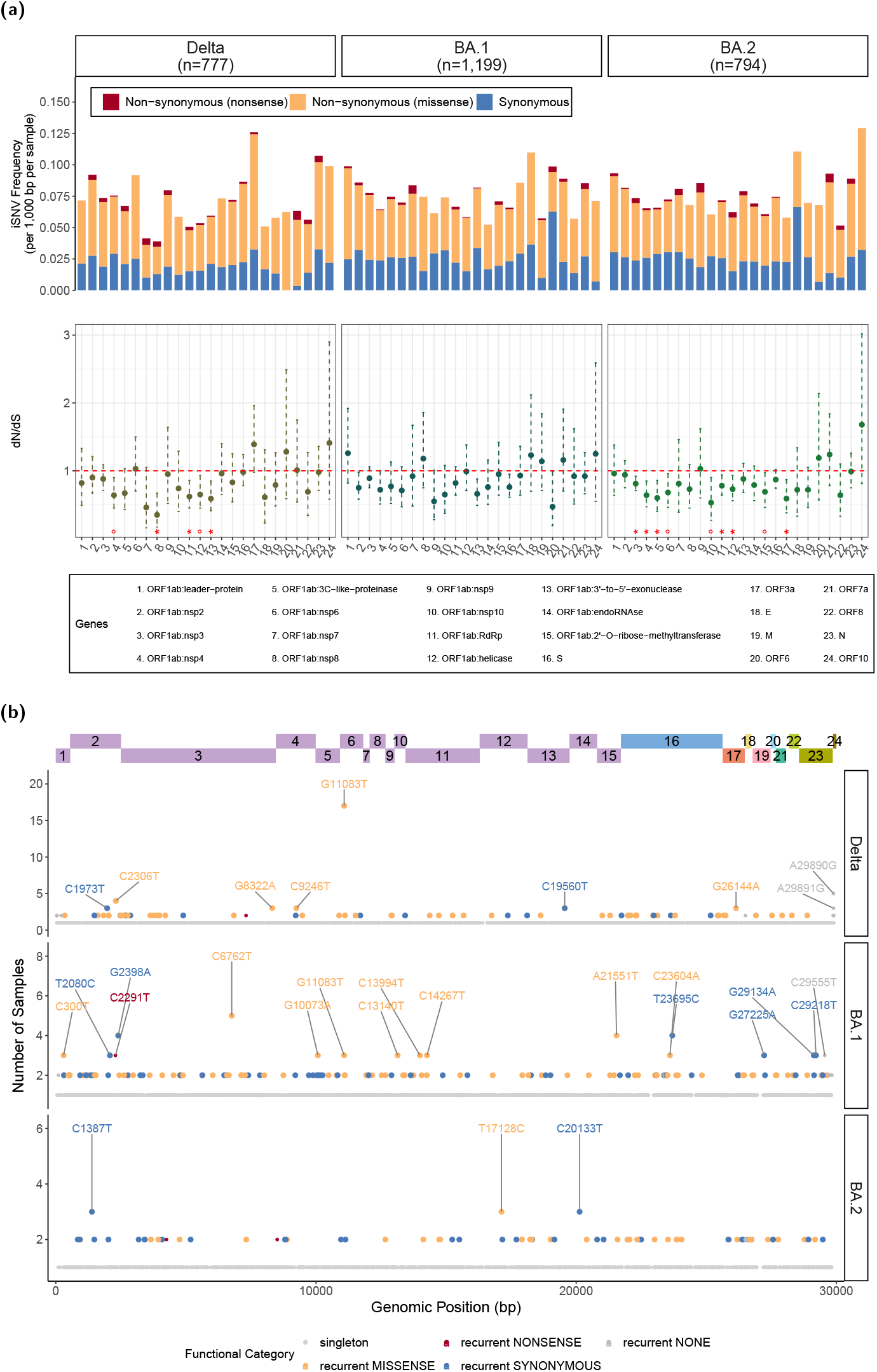
**(a) Frequency and dN/dS of iSNVs across genes in each VoC**. The stars at the bottom of the the dN/dS plot indicate the significance level, after Benjamini-Hochberg correction for multiple testing. **(b) Distribution of recurrent iSNVs across the genome**. Recurrent iSNVs, are defined as iSNVs present in 2 or more samples per VoC. The y-axis scale differs across VoCs

There was consistent evidence of purifying selection on genes in the ORF1ab polyprotein, which encodes the viral replication machinery, with statistically significant low dN/dS values among Delta and BA.2 isolates after controlling for multiple testing with a Benjamini-Hochberg correction (Figure 3A). Nsp3-5 consistently showed purifying selection across all VoCs, with statistically significant low dN/dS values for BA.2 isolates and low but not statistically significant dN/dS values for Delta and BA.1. Nsp3 and nsp5 are both proteases which cleave the viral polyprotein, thereby enabling the release of functional non-structural proteins. [46] Nsnp 3 and Nsp4 are both involved in the assembly of vesicles in which viral replication takes place. [47] The genes encoding the RNA-dependent RNA polymerase (RdRp), helicase, and 3’ to 5’ exonuclease, all essential for viral replication, also displayed dN/dS values below 1, with significant evidence of purifying selection observed in Delta and BA.2. [48] Nsp8, a key cofactor for RdRp that primes RNA synthesis and enhances its processivity [48], was also observed to be under significant purifying selection among Delta isolates. These findings are consistent with previous reports of purifying selection on acting on ORF1ab as a whole, and specifically on most of the same genes we identified: namely nsp2-4, nsp7-8, as well as the RdRp, helicase and exonuclease. [6] They suggests that these ORF1ab proteins, notably the viral proteases and replication machinery, are highly conserved due to their essential roles in maintaining efficient viral replication within hosts. Although statistical significance was not reached in other ORF1ab genes, most had dN/dS ratios below 1, suggesting that they are also subject purifying selection, albeit under weaker selective pressure or with insufficient variation to reach significance, possibly due to their limited size. In BA.2, we also found significant evidence of purifying selection acting on ORF3a, an accessory protein that forms ion channels in host cell membranes, thereby promoting virion release and inducing host cell death. Though ORF3a is not essential for viral replication or assembly, it modulates viral-host interaction, which may explain its selection.

No genes showed evidence of positive selection. Notably, the spike (S) and nucleocapsid (N) genes consistently showed dN/dS values close to 1 in all three VoCs, with no evidence of significant selective pressure after multiple testing adjustment (Figure 3A). Although these findings are somewhat unexpected, given these proteins’ direct exposure to the immune system and the steadily growing state of population immunity through infections and vaccination over the course of the study, they are consistent with previous reports. [6]

The dN/dS patterns observed across genomic regions in each VoC may reflect their unique viral characteristics or the distinct immune landscapes they encountered, as suggested by previous reports. For instance, Gu et al. [29] found that ORF1ab was subject to strong purifying selection within hosts among BA.2 isolates in both vaccinated and unvaccinated individuals, although genes within ORF1ab were not analyzed individually. In contrast, they found that other genes experienced positive selection within some vaccinated hosts, depending on the VoC and vaccination regimen. For instance, ORF10 - which in our study consistently displayed high dN/dS without reaching statistical significance for positive selection - was commonly found under positive selection in vaccinated hosts. Interestingly, no genes were found to be under either selection of either sort within hosts among BA.1 isolates. The BA.1 wave was extremely rapid, replacing Delta within weeks and peaking roughly one month after introduction before itself being displaced by BA.2 (Figure S3). Such rapid exponential spread, driven by greater fitness, may have reduced the effective impact of purifying selection, which could influence the intrahost variation observed during this period.

#### Signals of Selection Among Recurrent iSNVs

Another signature of selection is convergent evolution, in which the same character state arises multiple times independently. To investigate evidence for this in our dataset, we identified 550 distinct recurrent intra-host mutations, defined as iSNVs appearing in 2 or more isolates not linked by transmission (Figure 3B). Among these, 30 occurred in non-coding regions, while the remainder included 173 synonymous and 347 non-synonymous mutations. The ratio of non-synonymous-to-synonymous mutations (N/S) for recurrent iSNVs did not differ significantly from that of singleton iSNVs (2.01 vs 2.22; chi-square p-value=0.33; Table S1). This is consistent with previous reports that found no evidence of positive selection among recurrent iSNVs, which might have suggested sites of convergent evolution. [8]

Focusing on specific VoCs, we identified 79 distinct recurrent mutations among iSNVs in Delta, 141 in BA.1, and 72 in BA.2 isolates (Figure S9b). In BA.2, non-synonymous mutations were significantly enriched among singleton iSNVs relative to recurrent ones (chi-square p=0.03, Table S1)), indicating stronger purifying selection acting on recurrent iSNVs. In contrast, Delta and BA.1 showed no significant enrichment of non synonymous mutations in recurrent iSNVs compared with singletons (chi-square p=0.40 and chi-square p=0.25; Table S1), which may reflect either similarly weak purifying selection acting on both singletons and recurrent iSNVs, or slightly stronger purifying selection on recurrent mutations, with positive selection driving convergent evolution at a few sites. Notably, we identified one highly recurrent Delta iSNV (G11083T), possibly reflecting positive selection at this site, as detailed below.

#### A Potential Site of Positive Selection

The most recurrent iSNVs in our dataset, G11083T, was detected in 17 Delta isolates (2.2%) and 3 BA.1 samples with consistently high AAFs (median 0.87) (Figure 3B, Figure S11). This recurrent missense mutation results in a leucine-to-phenylalanine substitution at position 37 (L37F) of NSP6, a transmembrane viral protein that likely plays an important role in both virus replication efficiently and innate immune evasion. [47, 49] Specifically, NSP6 is involved in remodeling host cell membranes to form replication vesicles, and restricts the expansion of autophagosomes, thereby limiting the host cell’s ability to degrade viral components. The L37F mutation has previously been reported as an intra-host variant [7, 45] and a homoplasic consensus site, which has arisen multiple times across clades and geographies in the GISAID repository while not becoming fixed in any lineage (Figure S11). [7, 45, 50] Prior epidemiological studies have associated it with milder or asymptomatic infections. [51, 52] Although the exact functional impact of L37F remains to be experimentally validated, bioinformatic analyses suggest potential fitness advantages related to virus-host interactions. Specifically, L37F may disrupt interferon signaling pathways and modulate NSP6-mediated autophagy, thereby enhancing viral replication and promoting immune evasion. [51] Consistent with this, Delta isolates carrying G11083T in our study exhibited significantly lower Ct values compared to those without the mutation (median: 19.6 vs 27.8; Wilcoxon p = 6.9 *×* 10^−5^, Figure S12), indicating higher viral loads.

The repeated emergence of L37F across different genomic backgrounds in the global phylogeny and in our dataset indicates it is frequently acquired yet does not persist in the population, which suggests a fitness trade-off in which enhanced adaptation within individual hosts may come at the expense of efficient transmission. However, our finding that infections with this iSNV have significantly higher viral loads is then difficult to explain since existing evidence associates higher viral loads with an increased risk of transmission. [53] Indeed, we and others have found that iSNVs are very rarely shared through transmission, only being found 9 of 3,919 potential transmission events. Yet in two of those 9 cases the shared iSNV is G11083T, suggesting that this mutation can not only be transmitted, it is much more likely to be transmitted. An alternative explanation is that the frequency of G11083T fluctuates during the course of infection, potentially driven by frequent back mutations or tissue tropism. Supporting this, G11083T has been observed as an iSNV in one anatomical niche but not another within the same patient, and transient variants have been documented in longitudinal samples from a chronic infection. [45, 5]

## CONCLUSIONS

In this study, we analyzed SARS-CoV-2 intra-host variation using genomic data generated through BU’s testing mandate. We observed limited, rarely transmitted intra-host diversity, suggestive of a narrow transmission bottleneck and indicating that shared iSNVs are highly specific but insensitive markers of transmission. Our findings further suggest that the emergence of novel VoCs with drastic antigenic shift is more likely to result from prolonged evolution in chronic infections or animal reservoirs, rather than during acute infections. We found evidence that incomplete purifying selection shapes within-host evolution, with selection pressures varying across VoCs. Overall, recurrent iSNVs were consistent with random mutation rather than mutational hotspots, though one (G11083T) may reflect positive selection. A key strength of our study is the stringent filtering used to address systematic bias, including the batch-based approach we developed to reduce contamination-derived false-positive iSNVs in a data-driven manner given the absence of technical sequencing replicates. Although further validation is needed to fully assess the sensitivity and specificity of these methods, their application allowed us to recover findings consistent with those from carefully controlled experimental studies. This highlights the value of rigorously filtered surveillance genomic data for investigating within-host viral variation, and the need for robust, data-driven approaches to minimize false-positive signals.

## Ethical approval

The BU Charles River Campus Institutional Review Board reviewed and approved the sequencing protocol (5693E).

## Author contributions

Data collection and sequencing: J.H.C and J.T. Conceptualization: L.C., B.P.T, W.P.H. Data Analysis: L.C. Investigation: L.C., B.P.T, W.P.H., J.H.C, B.S. Writing: L.C. wrote the original draft. All authors contributed to review and final proofreading.

