## Supplementary Material for "SARS-CoV-2 Intra-host Variation Shows Evidence of Transmission and Convergent Evolution in a University Surveillance Cohort"

#### False-positive iSNVs from cross-contamination

Since technical sequencing replicates were not included during library preparation, additional filtering steps were necessary to exclude false-positive iSNVs resulting from systematic bias, notably well-to-well (or "cross-sample") contamination. We first examined the patterns of contamination, then developed a heuristic to remove likely false-positive iSNVs from contamination.

##### Cross-sample contamination patterns

We examined recurrent iSNVs ($0.03 \leq AAF < 0.95$) within sequencing batches and found that many corresponded to lineage-defining mutations. We further observed that isolates belonging to the corresponding lineage generally still carried the mutation at major allele frequencies ($0.5 \leq AAF$), whereas isolates from other lineages exhibited the mutation at minor allele frequencies (AAF $<$ 0.5), suggesting that the AAF of a falsepositive iSNV depends on the allele present at that genomic position in the contaminated sample. This pattern was most noticeable for VoC-defining mutations on plates containing isolates belonging to different VoCs, but also observed across sublineages (See Example in Figure S4). We additionally found no evidence of physical linkage among falsepositive iSNVs, as isolates rarely harbored multiple associated lineagedefining mutations as iSNVs. This suggests that contamination mostly occurs during library preparation, when aerosolized amplicons can transfer between wells during PCR amplification. [1] Overall, we concluded that contaminating amplicons generate falsepositive iSNVs with minor allele frequency (AAF $<$ 0.5) when the original sample has the Wuhan reference allele at that position, and with major allele frequency (AAF  0.5) when the original sample carries a nonreference fixed SNP (as illustrated in fig. S5).

##### Heuristic for removing false-positive iSNVs from contamination

Based on our empirical observations, we established the following two-step heuristic:

1. **Remove false-positive iSNVs due to contamination as described in scenario #1 in Figure S5:** A minor iSNV (AAF $<$ 50%) was considered a contamination-derived false positive if:

   (a) The same mutation was detected as a major variant (AAF $\geq$ 50%) in at least one other sample from the same library plate (indicating a potential source of contamination); **and**

   (b) The mutation corresponded to a lineage-defining SNP (according to a mutation-annotated phylogenetic tree), suggesting that it was unlikely to be a true de novo iSNV.

   When these conditions were satisfied, the minor iSNV was removed from the dataset. Additionally, for any corresponding major iSNVs (50% $\leq$ AAF $<$ 95%) found in isolates on the same library plate, the AAF was restored to 1 to reflect fixation, thus accounting for the bidirectional nature of contamination (see scenario #1 in Figure S5).

2. **Removing false-positive iSNVs due to contamination as described in scenario #2 in Figure S5:**
   Any remaining major iSNV (50% $\leq$ AAF $<$ 95%) was considered a false positive resulting from scenario #2 in Figure S5 if:

   (a) The isolate did not carry any other iSNV at the same position (i.e., the major iSNV was not due to a true positive minor variant); **and**

(b) At least one other sample from the same library plate harbored the Wuhan reference allele as the major allele at that position (indicating a potential source of contamination); **and**

(c) The mutation corresponded to a lineage-defining SNP (according to a mutation-annotated phylogenetic tree), suggesting that it was unlikely to be a true de novo iSNV.

When these criteria were met, the AAF of this major iSNV was restored to 1 to reflect that it was in fact a fixed mutation.

#### Limitations

Although our heuristic is informed by careful review of empirical evidence, it has limitations, and relies on conservative assumptions:

- We remove minor false-positive iSNVs and restore the AAFs of major false-positive iSNVs to a fixed value, thereby making assumptions about the original allele in corresponding isolates. Although this approach reflects general empirical patterns, the 50% threshold to distinguish minor and major variants could result in misclassification of false-positive iSNVs with AAFs close to this cutoff, or severe contamination, may violate this assumption.
- In criteria 2a, major iSNVs present with a Wuhan reference minor allele are considered false-positive iSNVs, while those that carry a different minor allele are not. This approach presumes that low-frequency reversions to the reference allele among iSNVs are rare and less likely than contamination.
- In criteria 1b and 2c, we assume that any intrahost mutation corresponding to a lineage-defining SNP and shared between samples on the same library plate is more likely due to contamination than transmission linkage, regardless of the consensus sequence distance between those samples. Therefore, this approach inevitably removes true shared iSNVs between transmision pairs sequenced on the same plate, as these are indistinguishable from cross-contamination and are likely to arise due to the temporal proximity of samples linked by transmission.

Any violation of these assumptions would remove true biological signals, making our decontamination approach conservative: highly sensitive to false positives but potentially low in relative specificity.

In addition, the sensitivity of our method depends on the assumption that most contamination occurs at the library plate stage, rather than during other steps of the sequencing workflow. Although our empirical data predominantly showed contamination confined to individual library plates, rather than across aliquot plates, we cannot rule out the possibility of cross-sample contamination occurring at other stages. The sensitivity is also influenced by the completeness of the dataset. For instance, if a low-quality sample carrying the contaminating allele was discarded, it could not be used to identify and exclude associated contaminated iSNVs, potentially leading to residual contamination.

### False-positive iSNVs from residual sequencing bias

Upon removing likely false-positive iSNVs from contamination as above, most top recurring iSNVs exhibited patterns consistent with residual sequencing artifacts. For instance, specific mutations recurred across many isolates and sequencing batches at very low allele frequencies, rarely surpassing 0.05 (See example in Figure S6), and other mutations were highly recurrent within individual sequencing batches. (Figure S6) Overall, iSNVs recurred at low frequencies in our dataset, and the mean recurrence level of iSNVs (i.e. number isolates carrying it) decreased as we increased the AAF threshold for filtering iSNVs from 0.01 to 0.05 (AF>=0.01: 4.43, AF>=0.02: 2.18, AF>=0.03: 1.56, AF>=0.04: 1.46, AF>=0.05: 1.46). This is consistent with low-frequency iSNVs being indistinguishable from noise and largely not concordant between replicates. [2, 3] Before raising the AAF threshold, we used information from low-frequency iSNVs to identify additional potential false-positive iSNVs that may not be eliminated by AAF-based filtering alone. As such, we removed any minor allele iSNVs (AAF<0.5, n=35,708) corresponding to mutations that recurred at low frequency (below 0.05) in isolates across 5 or more different sequencing batches. In addition, we remove minor iSNVs (0.01<=AF<0.05) recurring within a batch more than twice, since we assume this sharing of minor alleles is more likely an artifact rather than convergent evolution or a transmission cluster (n=122,640). We also restore major iSNVS recurring within a batch more than twice, including at least once at fixed frequency (n=186). Upon this filtering step, the average iSNVs recurrence level is more stable across AF thresholds (AF>=0.01: 1.28, AF>=0.02: 1.15, AF>=0.03: 1.12, AF>=0.04: 1.11, AF>=0.05: 1.11). After this, only iSNVs with AAF >=0.03 are included in the final analysis. [2, 3]

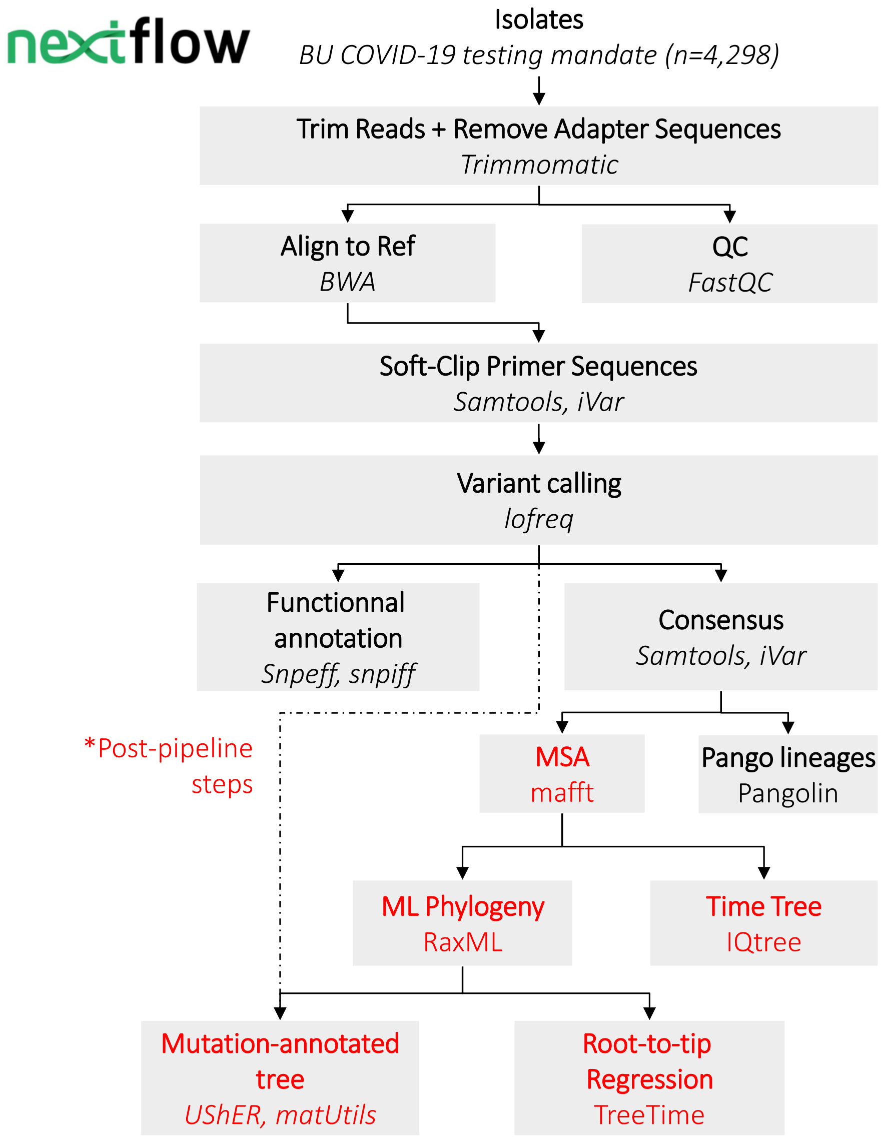


**Figure S1. Flowchart of the bioinformatic processing steps.** The code is for these processing steps is publically available on GitHub at: https://github.com/Leacavalli/Sars-cov-2-Intrahost-Variation.

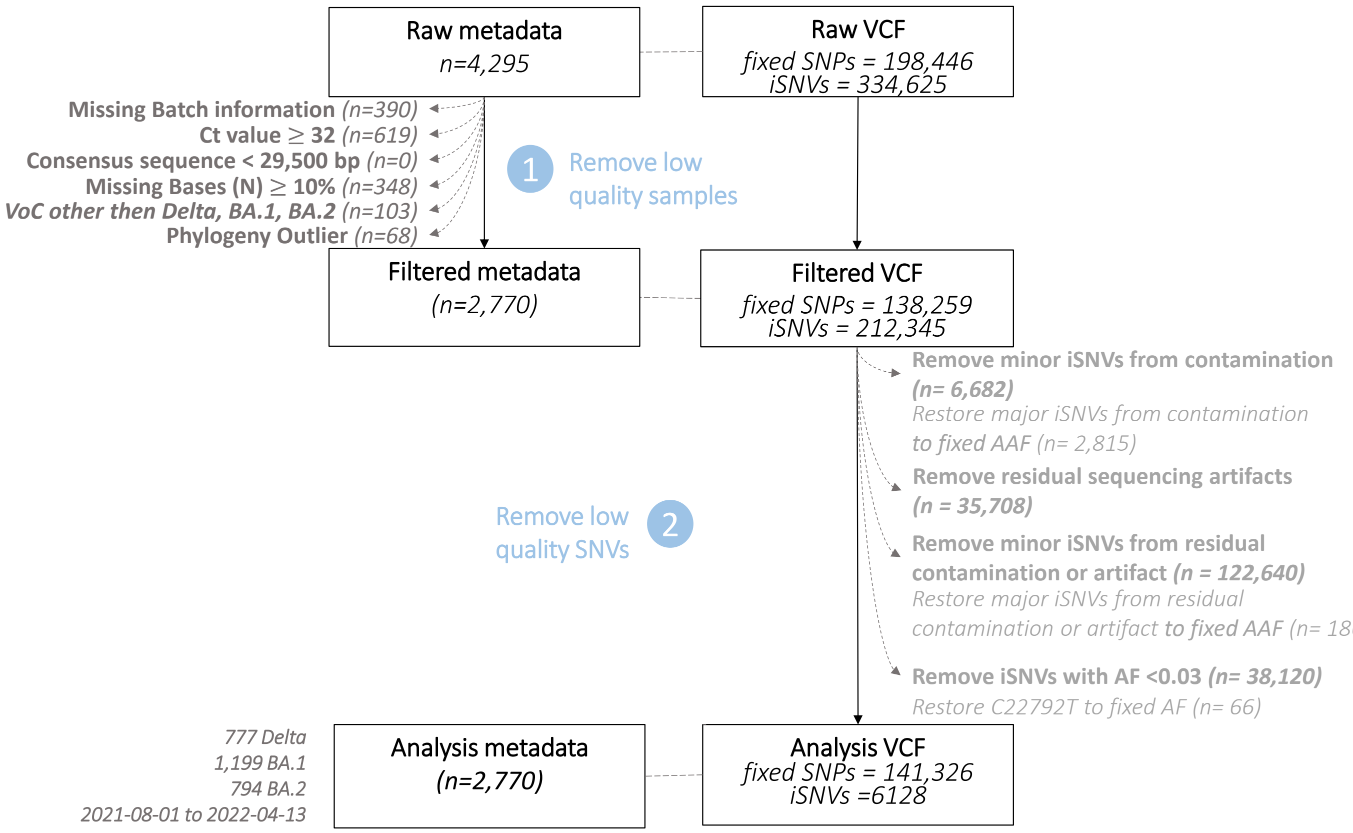


**Figure S2. Flowchart of the data filtering and cleaning process.** The metadata files are depicted on the left, while the multi-sample VCF file containing single nucleotides variant calls for all our isolates is shown on the right. The data cleaning process occurs in two stages: (1) The exclusion of low quality samples, and (2) The exclusion of low quality and false-positive iSNVs.

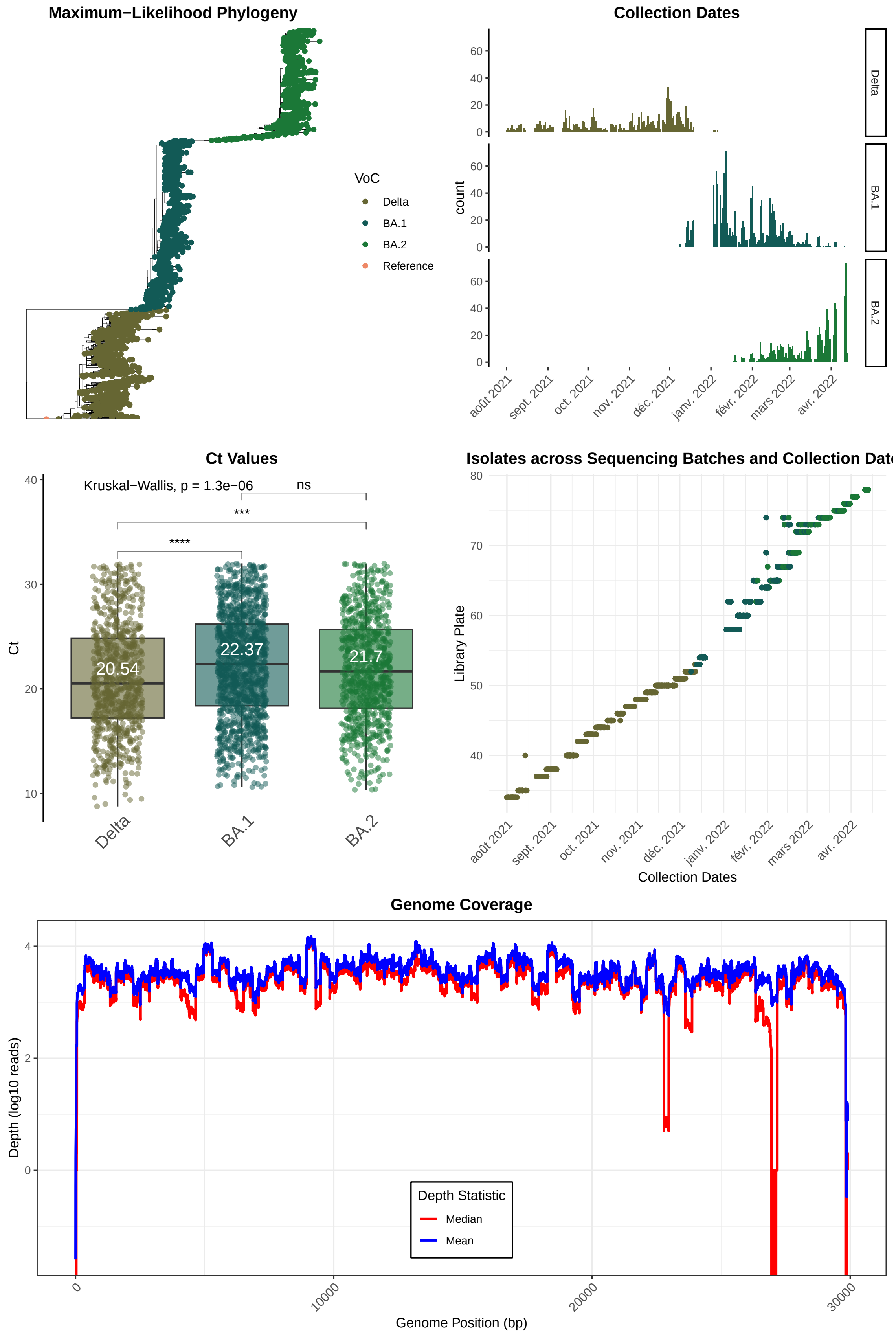


**Figure S3. Sample characteristics.** Consistent with previous reports, Delta isolates exhibit significantly smaller Ct values compared with BA.1 and BA.2 isolates, indicating higher viral loads.

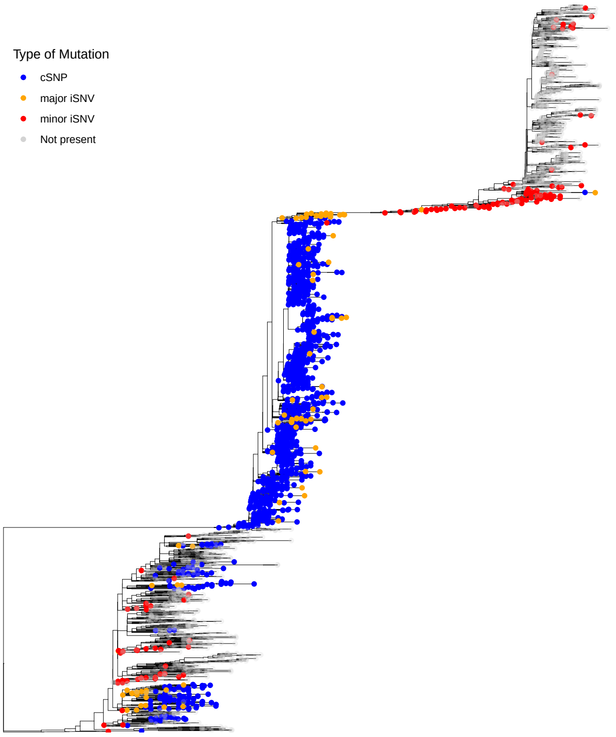


**(a)** Phylogenetic distribution of C21846T.

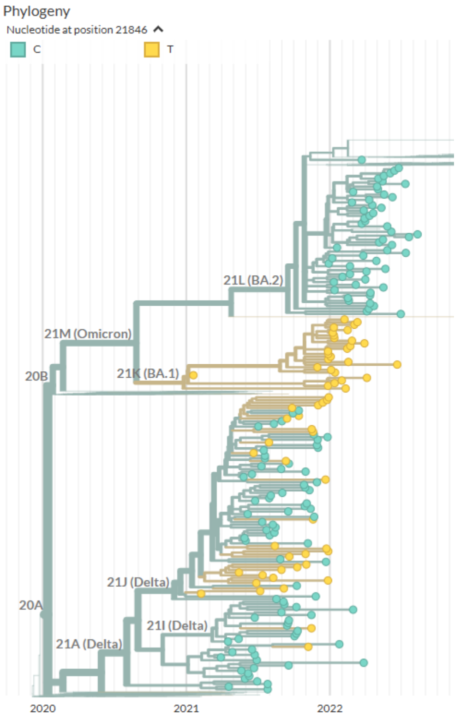


**(b)** Consensus allele at position 21,846 in GISAID (Nextstrain screenshot).

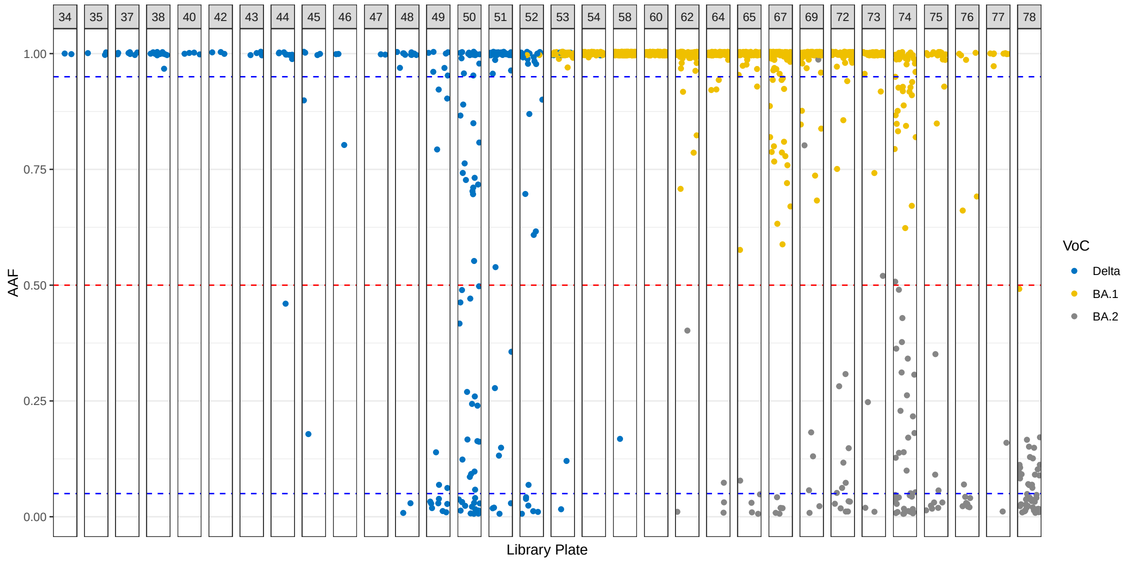


**(c)** Allele frequency (AF) of C21846T across sequencing batches.

**Figure S4. C21846T - an example of well-to-well contamination in our dataset.** C21846T is a BA.1-defining SNP, and is also present at consensus frequencies in some Delta sublineages. In our dataset, C21846T frequently appears as an iSNV in BA.2 isolates and Delta sublineages that do not typically carry the mutation, at minor allele frequency (AAF<0.5). The iSNV-carrying isolates are consistently found on plates with consensus-SNP carrying isolates (≥0.5), suggesting that C21846T is likely a false positive iSNV caused by cross-sample contamination.

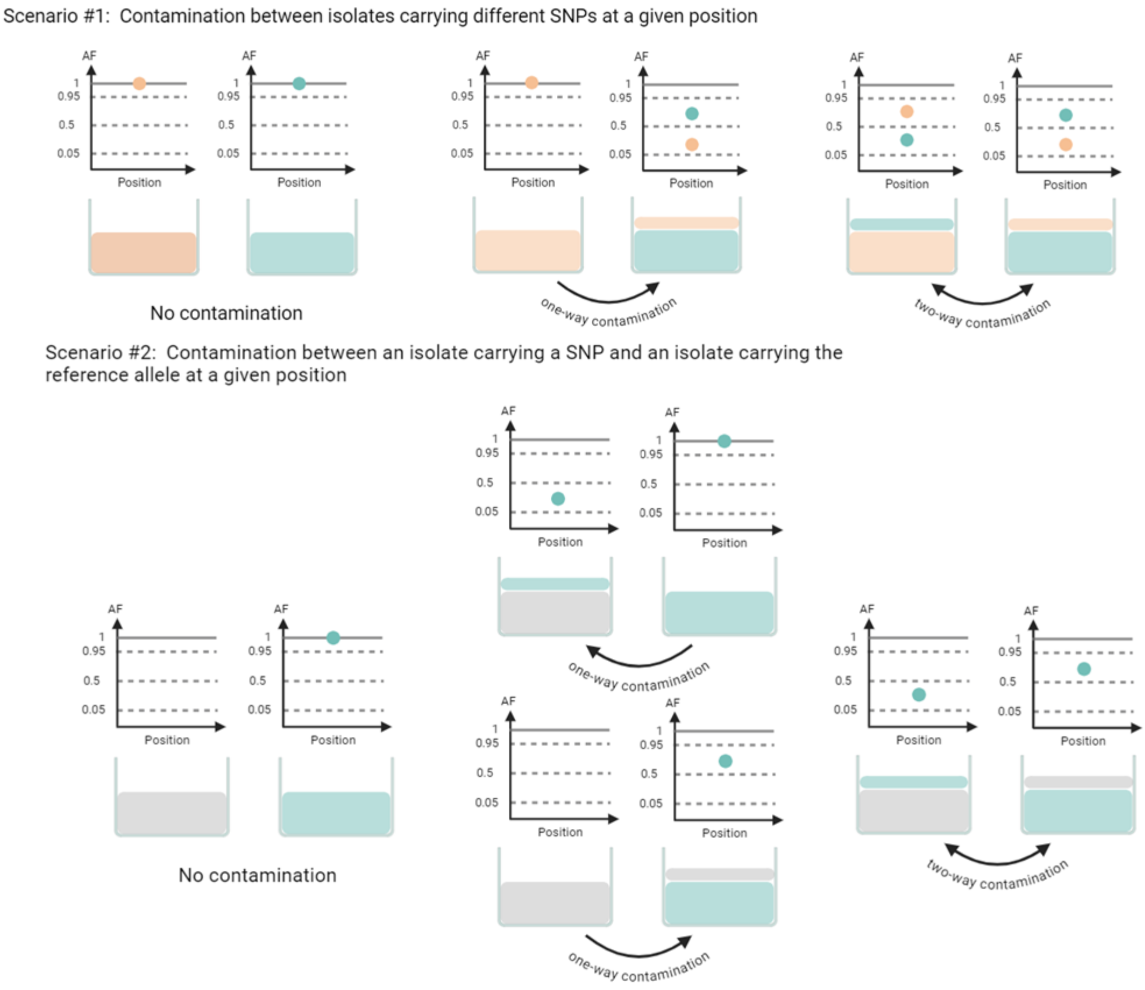


**Figure S5.** **Illustration of Possible Contamination Scenarios.**

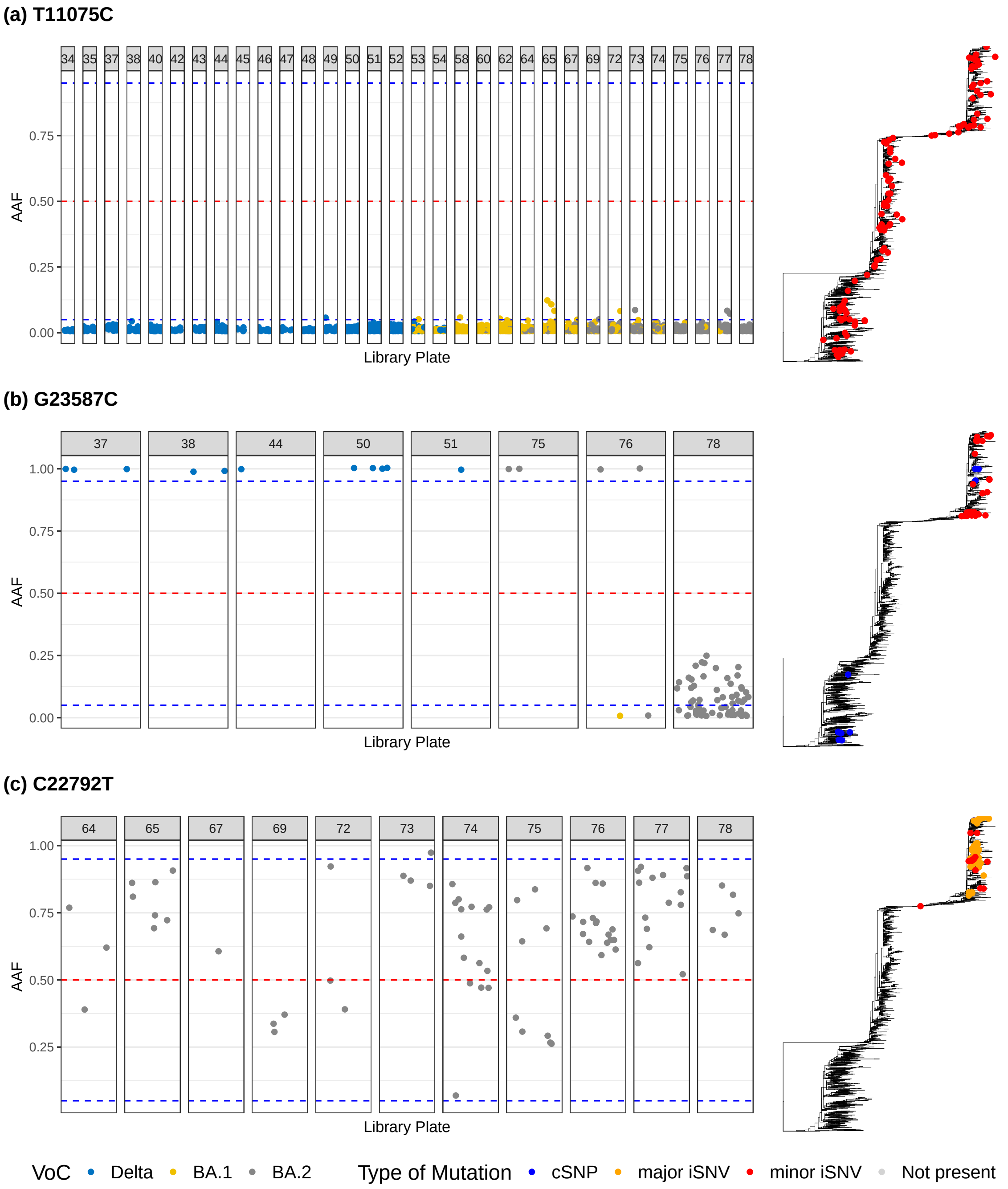


**Figure S6. Examples of false-positive iSNVs from residual bias upon minimizing contamination.** (a) T11075C is a previously reported residual systematic bias from sequencing artifact, appearing at low frequencies in many distantly-related isolates in our dataset (n = 2,138). [4] (b) G23587C represents likely residual contamination or artifact within a specific sequencing batch, recurrently detected at subconsensus frequencies in isolates (n = 63/130) processed on library plate 78. (c) C22792T is a known systematic artifact caused by failed primer binding. While it is a lineage-defining SNP for several BA.2 sublineages, its allele frequency remains consistently below the fixation threshold of 0.95.

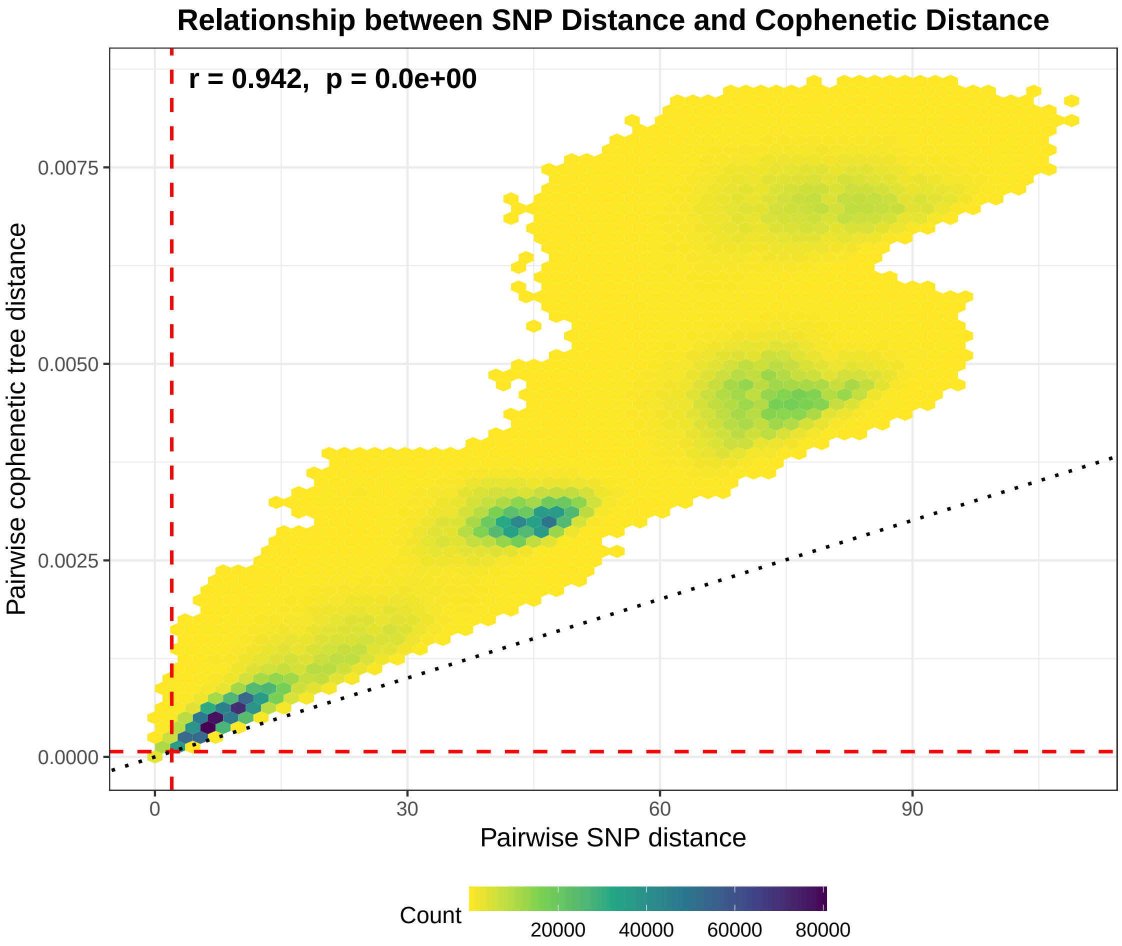


**Figure S7. Correlation between SNP differences and patristic distances.** SNP differences were counted from the multiple sequence alignment using SNP-dist [5]. Patristic distances were calculated from a maximum-likelihood phylogeny constructed with RAxML under the GTR+Gamma model, using the `cophenetic.phylo` function from the `ape` R package. Although the two measures are significantly correlated, SNP counts are systematically lower, likely because SNP differences do not account for site-specific variation in evolutionary rates and may therefore underestimate divergence times [6, 7]. Using cophenetic distances instead of SNP differences to identify potential transmission pairs is both more computationally intensive and more conservative, as indicated by the red dashed lines, but this approach makes fuller use of the sequence alignment.

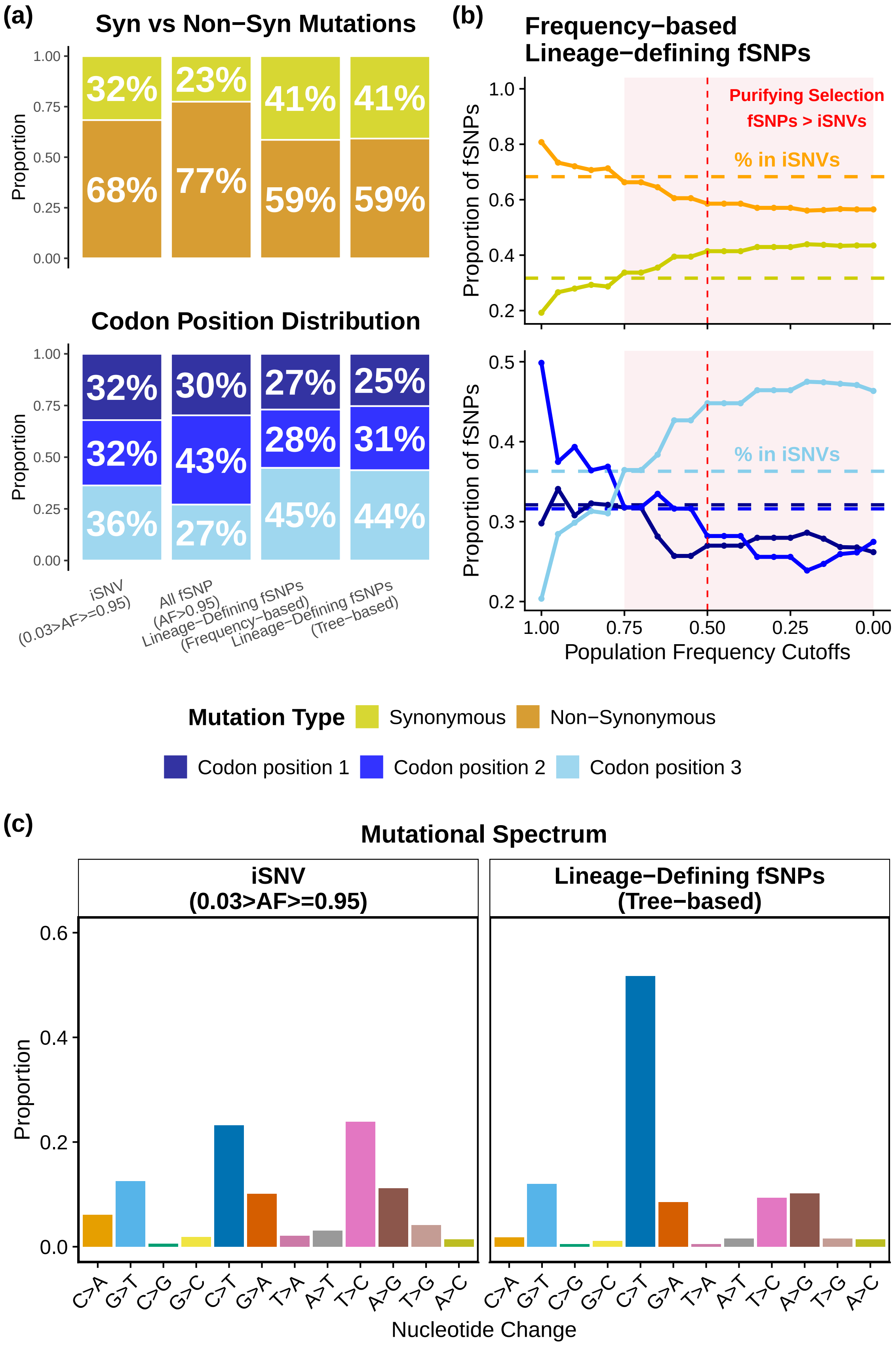
(a)
Syn vs Non−Syn Mutations
Proportion
32%
68%
23%
77%
41%
59%
41%
59%
Codon Position Distribution
32%
32%
36%
30%
43%
27%
27%
28%
45%
25%
31%
44%
iSNV (0.03>AF>=0.95)
All fSNP (AF>0.95)
Lineage−Defining fSNPs (Frequency−based)
Lineage−Defining fSNPs (Tree−based)
(b)
Frequency−based Lineage−defining fSNPs
Proportion of fSNPs
Purifying Selection
fSNPs > iSNVs
% in iSNVs
Population Frequency Cutoffs
Mutation Type
Synonymous
Non−Synonymous
Codon position 1
Codon position 2
Codon position 3
(c)
Mutational Spectrum
iSNV (0.03>AF>=0.95)
Lineage−Defining fSNPs (Tree−based)
Proportion
Nucleotide Change
C>A G>T C>G G>C C>T G>A T>A A>T T>C A>G T>G A>C

**Figure S8. Mutational Characteristics of iSNVs compared to fSNPs.** (a) Proportion of synonymous and nonsynonymous mutations (top) and the distribution of mutations by codon position (bottom) among iSNVs ($3\% \leq \text{AAF} < 95\%$), all fixed SNPs (fSNPs, $\text{AAF} \geq 95\%$), and lineage-defining fixed SNPs. Counting all fixed SNPs across samples skews the mutational spectrum toward nonsynonymous and third-codon-position mutations, largely due to repeated counting of lineage-defining SNPs. To address this, lineage-defining SNPs can be counted only once, defined either by population frequency (frequency-based) or by mutation-annotated phylogenetic trees (tree-based). (b) Proportion of synonymous and nonsynonymous mutations (top) and distribution by codon position (bottom) across varying population frequency thresholds for defining frequency-based lineage-defining fSNPs. When the threshold is set at 75% or lower, the proportion of nonsynonymous and third-codon-position mutations among fSNPs falls below that of iSNVs, indicating stronger purifying selection. These proportions plateau at the 50% threshold, closely matching those obtained with the tree-based definition. (c) Mutational spectrum in iSNVs and tree-based lineage-defining fSNPs. Among fSNPs, we observe pronounced C>U over U>C and C>U over G>A asymmetries, consistent with the expected signature of host APOBEC-mediated cytidine deamination acting preferentially on the positive-sense viral RNA strand. In contrast, among iSNVs, we detect strong C>U over G>A and U>C over A>G asymmetries, indicating pronounced strand bias in mutation patterns but without a strong directional preference between C>U and U>C transitions. This pattern may reflect intrinsic features of the viral replication process, with iSNVs capturing the immediate outputs of viral polymerase errors or short-term editing activity, while fSNPs - having persisted over multiple replication and transmission cycles  may reflect the cumulative effects of prolonged exposure to APOBEC-mediated editing.

**Table S1.** Categories and summary statistics for singleton and recurrent iSNVs across Delta, BA.1, and BA.2 lineages.

| | Delta (n=777) | BA.1 (n=1,199) | BA.2 (n=794) | Overall (n=2,770) |
|---|---|---|---|---|
| **Singleton iSNVs** | | | | |
| Total iSNVs | 1,569 | 2,328 | 1,585 | 4,873 |
| Non-coding | 54 | 51 | 31 | 110 |
| Synonymous (S) | 434 | 743 | 494 | 1,481 |
| Non-synonymous (N) | 1,081 | 1,534 | 1,060 | 3,282 |
| N/S | 2.49 | 2.06 | 2.15 | 2.22 |
| $d_N/d_S$ (95% CI) | 0.79 (0.70–0.90) | 0.82 (0.74–0.90) | 0.84 (0.75–0.95) | 0.81 (0.76–0.87) |
| **Recurrent iSNVs** | | | | |
| Total iSNVs | 184 | 304 | 147 | 1,244 |
| Non-coding | 24 | 19 | 2 | 71 |
| Synonymous (S) | 34 | 108 | 66 | 398 |
| Non-synonymous (N) | 126 | 177 | 79 | 775 |
| N/S | 3.71 | 1.64 | 1.20 | 1.95 |
| $\chi^2$ p-value vs Singleton N/S | 0.06 | 0.09 | 0.001 | 0.07 |
| $d_N/d_S$ (95% CI) | 1.07 (0.71-1.61) | 0.74 (0.57-0.96) | 0.46 (0.32-0.66) | 0.78 (0.68-0.89) |
| Total unique mutations | 79 | 141 | 72 | 550 |
| Non-coding | 10 | 9 | 1 | 30 |
| Synonymous (S) | 16 | 50 | 32 | 173 |
| Non-synonymous (N) | 53 | 82 | 39 | 347 |
| N/S | 3.31 | 1.64 | 1.22 | 2.01 |
| $\chi^2$ p-value vs Singleton N/S | 0.40 | 0.25 | 0.03 | 0.33 |
| $d_N/d_S$ (95% CI) | 1.07 (0.58–1.96) | 0.75 (0.51–1.09) | 0.46 (0.27–0.78) | 0.81 (0.67–0.99) |

(a)

(b)

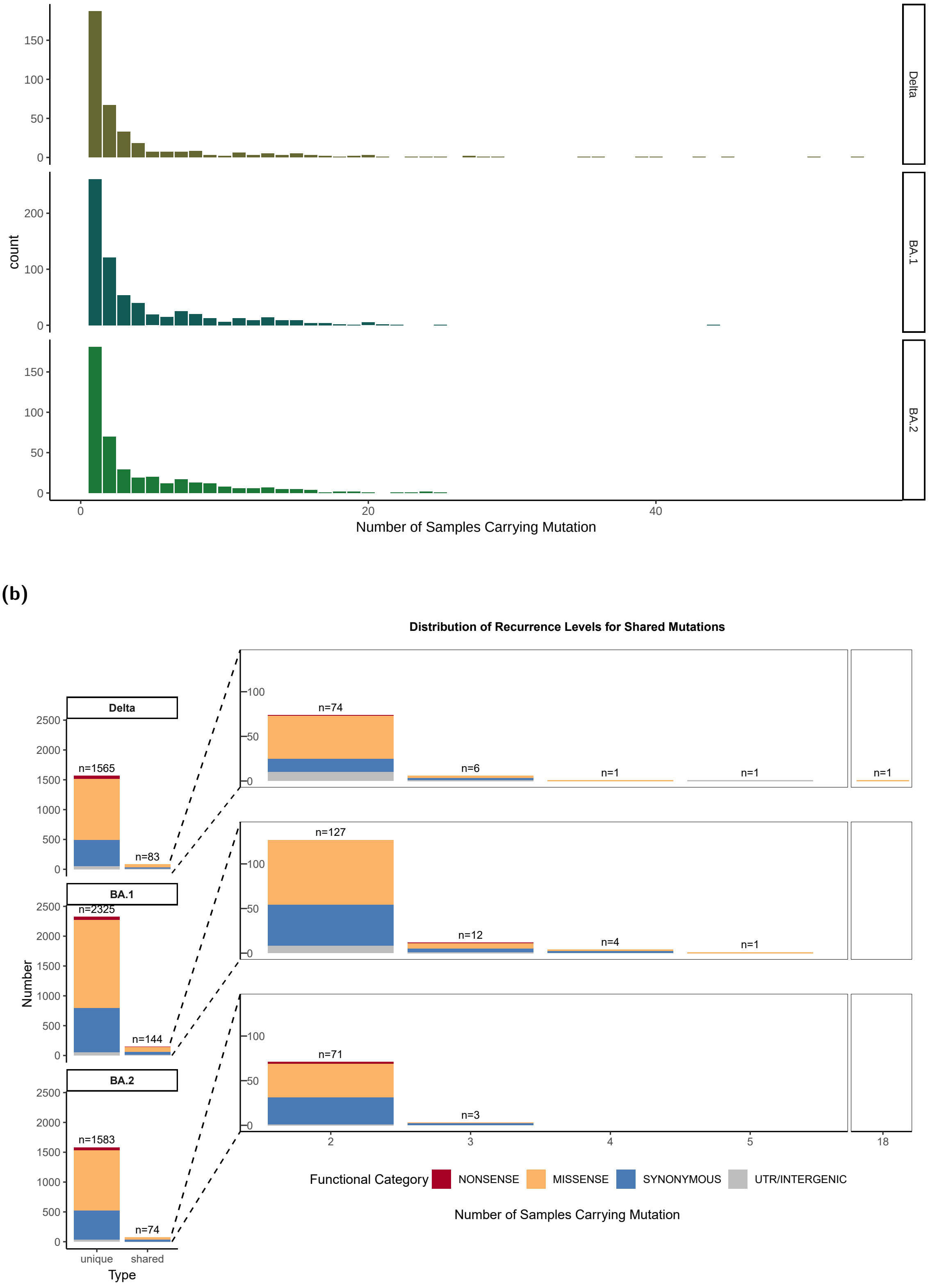


**Figure S9. (a) Distribution of the number of iSNVs per sample across VoCs**, and **(b) Counts of unique and shared iSNVs as well as their recurrence distribution across VoCs.** "Unique" mutations occur in a single sample, while "shared" mutations are found in multiple samples. The distribution of shared iSNVs by recurrence level (number of samples in which a shared iSNV appears) is also shown.

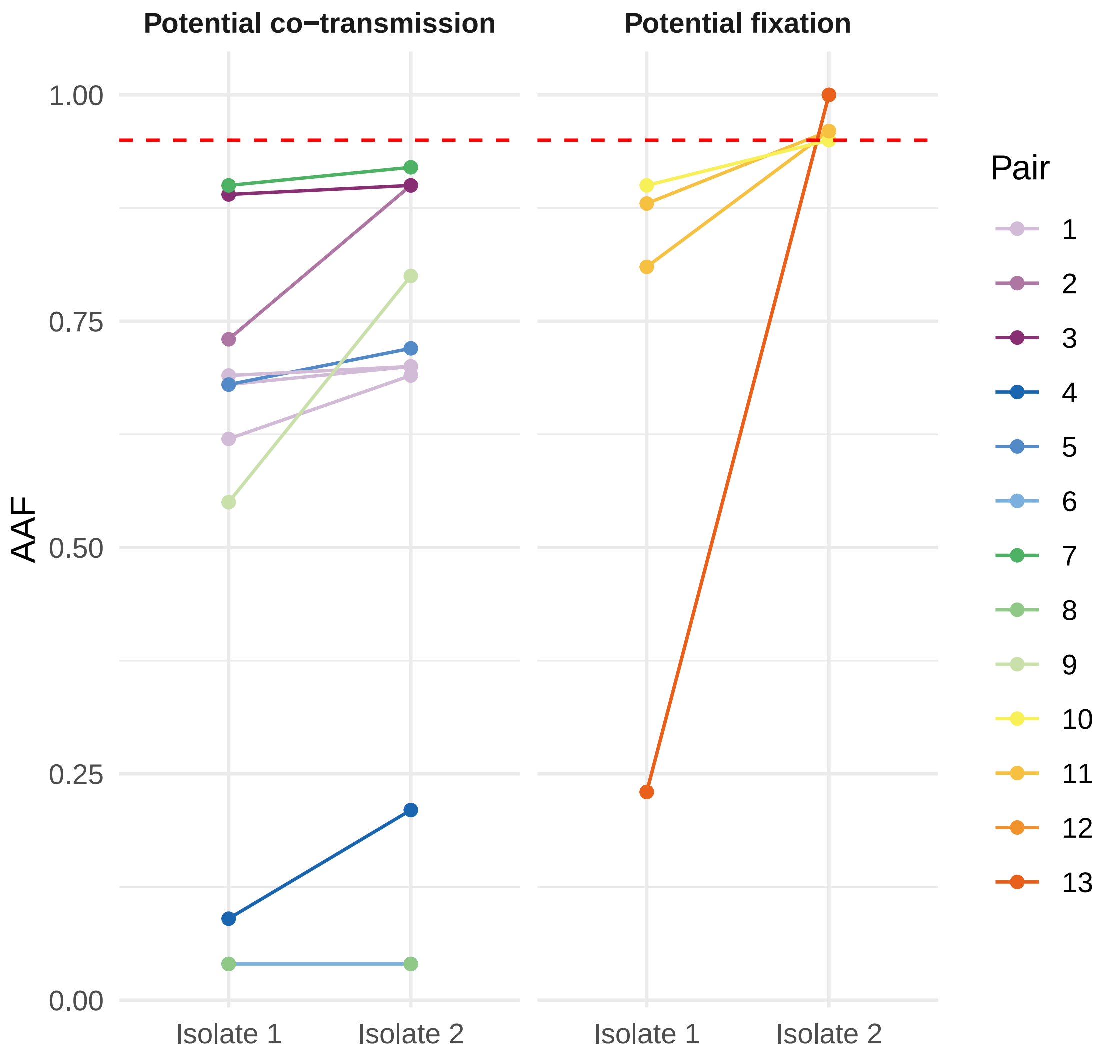


**Figure S10. Allele frequencies of shared iSNVs in potential transmission pairs.** Each line connects the alternative allele frequency (AAF) for a given iSNV between paired isolates. The left panel shows potential co-transmission events, where the iSNV is present in both isolates. The right panel shows potential fixation events, where the iSNV rises to fixation (AAF $\geq$0.95) in one of the isolates. Each pair is color-coded. The red dashed line marks the consensus threshold (AAF = 1).

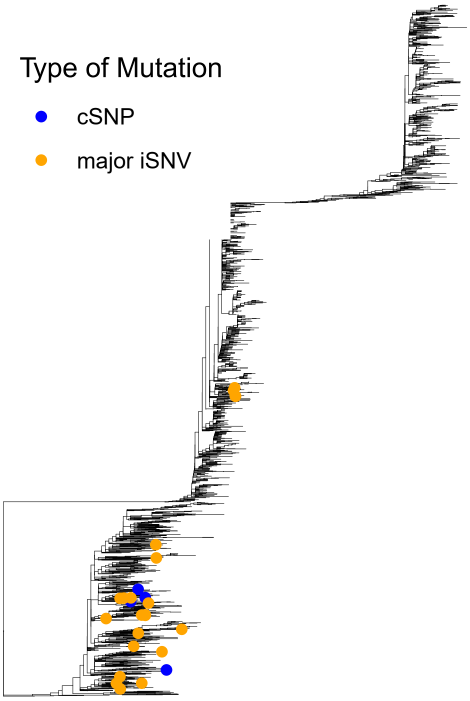


**(a)** Phylogenetic distribution of G11083T.

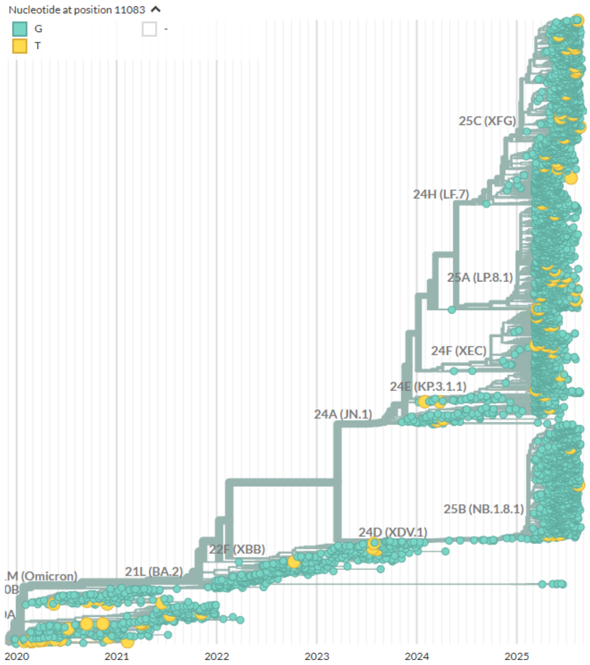


**(b)** Consensus allele at position 11,083 in GISAID (Nextstrain screenshot).

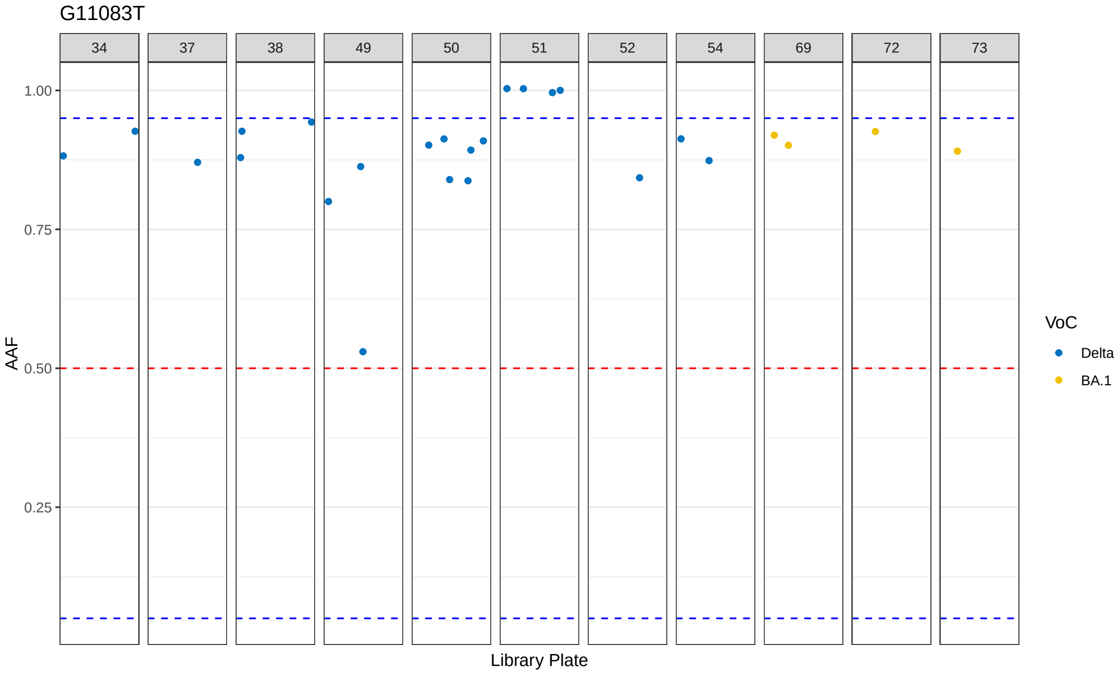


**(c)** Allele frequency (AF) of G11083T across sequencing batches.

**Figure S11. G11083T - a highly recurrent mutation among iSNVs.**

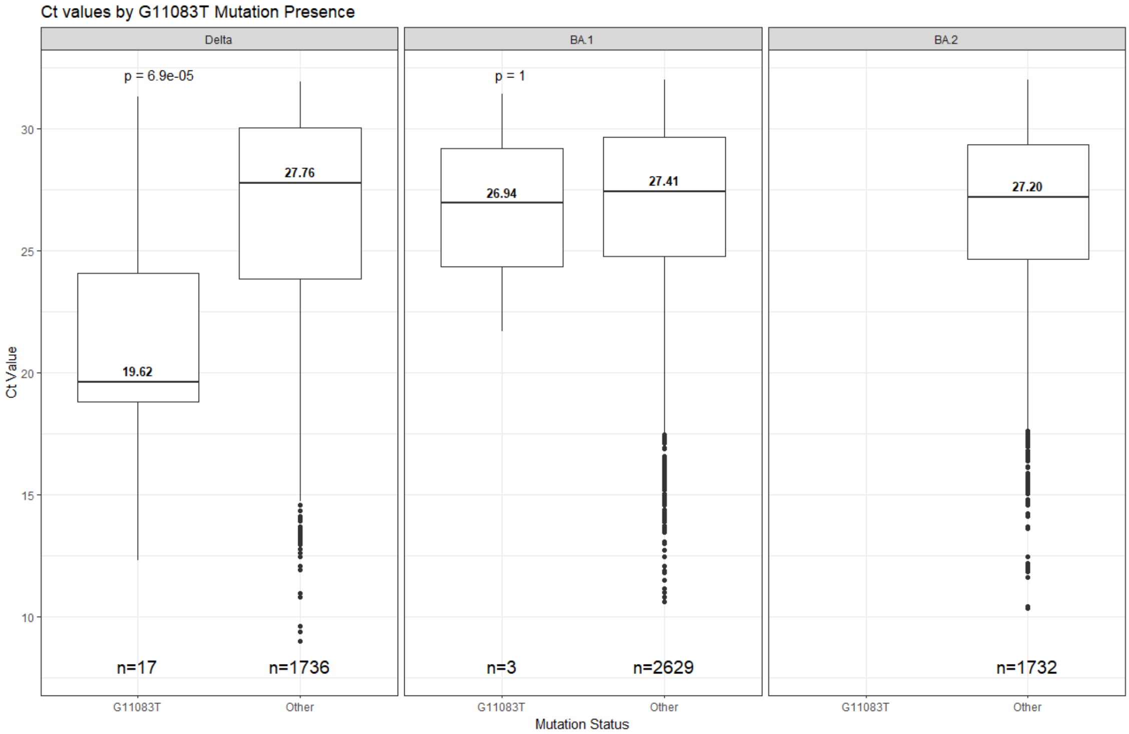


**Figure S12. Distribution of Ct values for isolates with and without the iSNV G11083T.**
